# Mutualistic rhizobia harbor genetic variation for traits related to parasite infection

**DOI:** 10.64898/2026.01.20.700730

**Authors:** Addison D Buxton-Martin, Corlett Wolfe Wood

## Abstract

Nutritional mutualisms are defined by resource exchange, but growing evidence suggests that these interactions also shape resistance to parasites and pathogens. The evolutionary significance of this phenomenon is unclear, primarily because research has focused on mutualist-mediated plasticity rather than on mutualist contributions to heritable variation in the outcome of infection. To test whether nutritional mutualists harbor genetic variation for infection-related traits, we performed a quantitative genetic experiment in the nutritional mutualism between legumes and nitrogen-fixing rhizobia. We paired 10 mutualistic *Sinorhizobium meliloti* rhizobia strains with 20 *Medicago truncatula* plant genotypes in an incomplete factorial design, and experimentally infected plants with parasitic root-knot nematodes (*Meloidogyne hapla*). We used this design to estimate plant and rhizobia contributions to genetic variation in four infection-related traits: resistance, tolerance, virulence, and mutualism robustness to infection. Rhizobia contributed directly to genetic variation in virulence, and to parasite resistance and mutualism robustness via genotype-by-genotype interactions. Rhizobia did not contribute to genetic variation in tolerance. The effect of rhizobia on parasite resistance was partially explained by their effect on root growth. Our results raise the possibility that some nutritional mutualists play a role comparable to defensive mutualists in shaping the evolutionary potential of host defense traits.

**Teaser text:** Nutritional mutualisms are everywhere. These mutualisms are defined by the exchange of resources between partners, but many have secondary effects on resistance to parasites, pathogens, or herbivores. Here, we show that a textbook nutritional mutualist (nitrogen-fixing rhizobia) harbors genetic variation for the host’s response to parasite infection. Our results imply that nutritional mutualists can impact the evolutionary potential of infection-related traits in their hosts, suggesting that these mutualisms may be overlooked drivers of defense evolution.

## Introduction

Mutualisms are traditionally categorized by the benefits they provide. Nutritional mutualists exchange resources, defensive mutualists provide protection, and transportation mutualists facilitate movement (Bronstein, 2015). Yet mutualist impacts on their hosts can extend well beyond the primary benefit of the interaction (Batstone et al., 2022; Friesen et al., 2011; Fronk & Sachs, 2022; Ma et al., 2020; O’Brien et al., 2021). Nutritional mutualisms, for example, frequently affect resistance to parasites, pathogens, and herbivores (Buxton-Martin et al., 2025; Koch & Schmid-Hempel, 2011; O’Keeffe et al., 2022). This observation indicates that effects on interactions with enemies are not limited to defensive mutualisms, and raises an important question: do non-defensive mutualisms also influence the evolution of defense traits?

There are now many examples of how nutritional mutualisms affect their host’s resistance to parasites, pathogens, or herbivores (reviewed in Wood et al., *in review*). This phenomenon is especially well studied in mycorrhizal fungi, which usually increase the host plant’s resistance to infection (Jung et al., 2012; Liu et al., 2023), but can decrease resistance as well (Eck et al., 2022; Miozzi et al., 2019). Nitrogen-fixing rhizobia increase their host plant’s vulnerability to a parasitic nematode (Buxton-Martin et al., 2025; Wood et al., 2018), as well as to herbivores (Simonsen & Stinchcombe, 2014). Similar effects on interactions with enemies have been documented in animals as well (Salem et al., 2015; Stevens et al., 2021). From these examples, it is clear that nutritional mutualists can have a profound impact on their host’s interactions with enemies. Furthermore, unlike defensive mutualists, nutritional mutualists are not always protective, implying that nutritional and defensive mutualisms may have markedly different consequences for the evolution of host traits related to defense.

In defensive mutualisms, symbionts contribute directly to the evolution of defense via heritable variation in host traits that resides in symbiont populations (reviewed in Vorburger & Perlman, 2018). In other words, defense is jointly determined by host and symbiont genomes. For example, *Drosophila* fruit flies inoculated with different *Wolbachia* strains differ in resistance to viral infection, indicating that *Wolbachia* harbors genetic variation for antiviral resistance (Martinez et al., 2014, 2017). In a landmark study, (Oliver et al., 2005) found that parasite resistance was explained by the genotype of the *Hamiltonella defensa* symbiont, not the aphid host. Subsequent studies, primarily in pea aphids (*Acyrthosiphon pisum*), established that differences in protection among defensive symbiont strains are common (Hafer-Hahmann & Vorburger, 2020; McLean et al., 2020; Parker et al., 2017; Rouchet & Vorburger, 2014; Schmid et al., 2012). In parallel, experimental (co)evolution, primarily in *Caenorhabditis elegans* nematodes, has directly linked defensive symbionts to the evolution of resistance and tolerance, as well as pathogen virulence (Ford et al., 2016, 2017; Hoang et al., 2024; King & Bonsall, 2017; Rafaluk-Mohr et al., 2022; Smith et al., 2025).

By contrast, the evolutionary significance of *non-defensive* mutualist effects on interactions with enemies remains largely untested. This is primarily because research in non-defensive mutualisms has emphasized mutualist-mediated *plasticity* over mutualist effects on *heritable variation* in infection-related traits. A few studies have reported differences in enemy interactions between hosts inoculated with different nutritional mutualist strains (Buxton-Martin et al., 2025; Simonsen & Stinchcombe, 2014). For example, *Medicago* plants inoculated with two different rhizobia strains differed in resistance to parasitic nematodes, as well as in their transcriptomic response to nematode infection (Buxton-Martin et al., 2025). However, two-strain experiments like this cannot directly estimate the mutualist’s contribution to genetic variation in infection-related traits, because genetic variance is a population-level property that requires many strains to measure accurately. Furthermore, benchmarking mutualist genetic variation against host genetic variation is essential to gauge the mutualist’s relative contribution to the evolutionary potential of a host trait. We’re aware of only one study in any mutualism that directly compared host and mutualist contributions to genetic variance in infection-related traits (Parker et al., 2017), in an aphid defensive symbiosis.

Here, we tested whether nutritional mutualists influence the evolutionary potential of host traits related to parasite infection. We asked three questions: First, do nutritional mutualists harbor genetic variation for infection-related host traits? Second, how much genetic variation do mutualists contribute to these traits, relative to hosts? Finally, do nutritional mutualists contribute equally to parasite resistance, tolerance, and virulence? We addressed these questions in the tripartite interaction between legumes, nitrogen-fixing rhizobia, and plant-parasitic root knot nematodes. *M. truncatula* is an annual legume native to the Mediterranean (Cook, 1999) and forms a mutualistic symbiosis with nitrogen-fixing rhizobia, such as *S. meliloti*. Rhizobia are housed within nodules that develop from the roots of their host plant where they trade with the host plant for carbon-containing compounds. The northern root-knot nematode (*Meloiogyne hapla*) is an obligate endoparasite that forms galls on plant roots, where it siphons nutrients from the host (M. G. K. Jones & Goto, 2011; Koltai et al., 2001). A previous study found that *M. truncatula* plants inoculated with two different *S. meliloti* strains differed in their resistance to nematode infection (Buxton-Martin et al., 2025).

We paired 20 *Medicago truncatula* plant accessions and 10 *Sinorhizobium meliloti* rhizobia strains in an incomplete factorial design, and infected half of the plants with parasitic nematodes (Figure 1A). We then partitioned total genetic variation in four infection-related traits (described below) into genetic variation in the host (direct genetic variance; G_Host_), genetic variation in the mutualism (indirect genetic variance; G_Rhizobia_), and genotype-by-genotype interactions between host and mutualist (intergenomic epistasis; G_Host × Rhizobia_) (De Lisle et al., 2022; Heath & Tiffin, 2007; Wade, 2007). Genotype-by-genotype interactions are pervasive in both nutritional and defensive mutualisms (Heath, 2010; Heath & Tiffin, 2007; Parker et al., 2017; Schmid et al., 2012), so it is crucial to account for genotype dependence when estimating genetic variation in joint host-symbiont traits.

**Figure 1.**
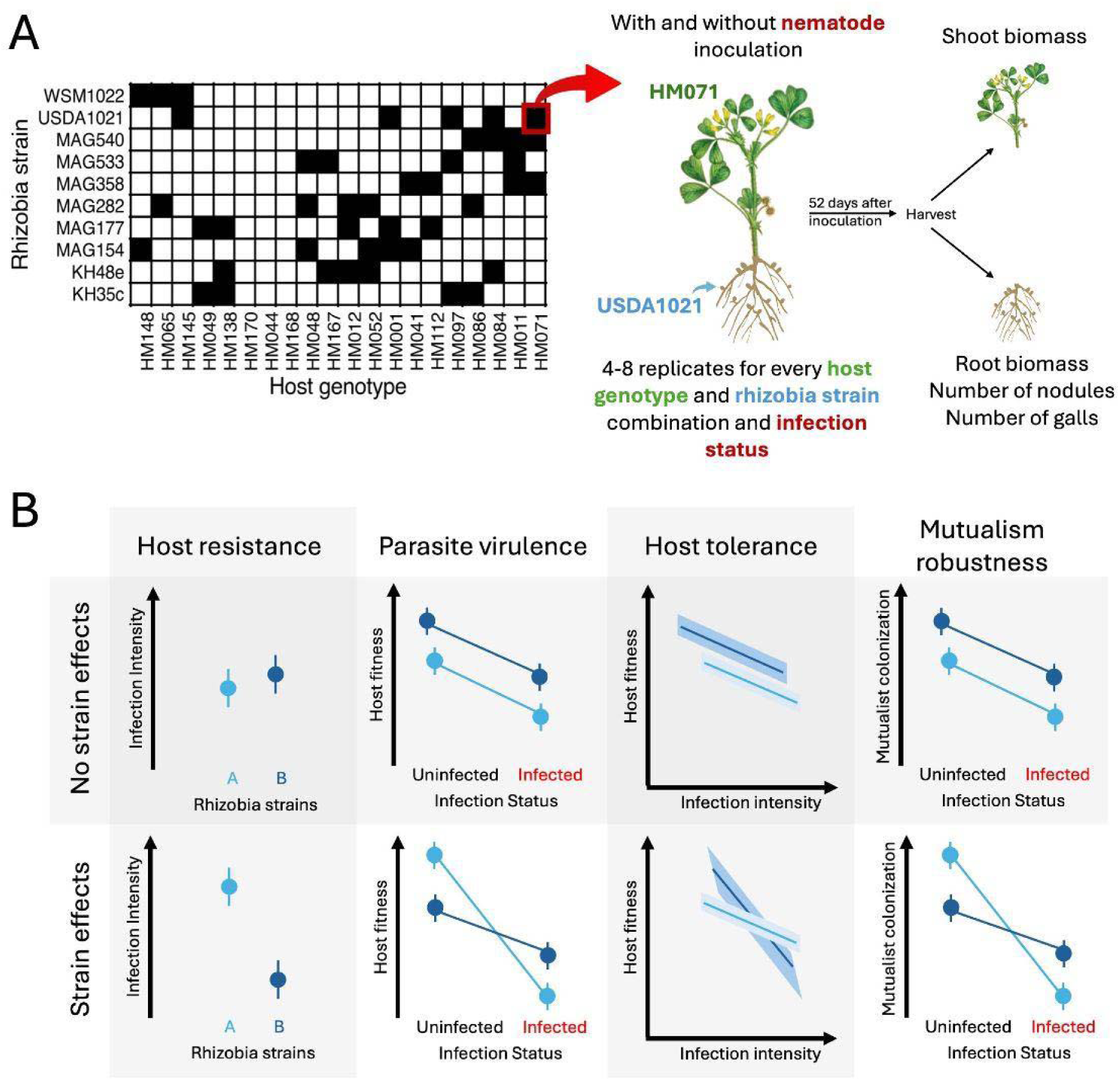
(A) A conceptual diagram showing our experimental design. (B) Conceptual diagrams that illustrate the absence (top row) and presence (bottom row) of rhizobia genetic variation for the four infection-related traits we measured. Rhizobia genetic variation for resistance is indicated by differences in infection intensity across rhizobia strains. Rhizobia genetic variation for virulence is indicated by a strain-dependent decrease in fitness in parasite-infected plants. Rhizobia genetic variation for tolerance is indicated by differences in the slope of the relationship between host fitness and infection intensity for different rhizobia strains. Rhizobia genetic variation for mutualism robustness is indicated by a strain-dependent decrease in mutualist colonization in parasite-infected plants.

We focused on four infection-related traits: resistance, tolerance, virulence, and mutualism robustness to infection (Figure 1B) (Bush et al., 1997). Resistance (the inverse of susceptibility) is the host’s ability to prevent parasite infection, and is measured by infection intensity (the total number of parasites on a host). Tolerance is the fitness cost per parasite, measured as the slope of the relationship between infection intensity and fitness. Virulence is the total fitness cost of infection, as measured as the difference in fitness between infected and uninfected hosts. All three of these traits are typically determined jointly by both the host’s and parasite’s genotype and differ in their implications for host-parasite interactions (Bruns et al., 2012; Jutzeler et al., 2024; Salvaudon et al., 2007). In addition to these three classic infection-related traits, we also measured mutualism robustness to parasite infection, defined as the impact of parasite infection on mutualist colonization (Figure 1B). This captures an often-overlooked cost of parasite infection: disrupted interactions with mutualists. Finally, we evaluated whether the rhizobial effect on infection intensity was related to their effect on host growth (i.e., nutritional mutualists increase plant size, and bigger plants host more parasites).

We found that rhizobia contributed directly to genetic variation in parasite virulence (G_Rhizobia_), and to parasite resistance and mutualism robustness via genotype-by-genotype interactions (G_Host x Rhizobia_). A path analysis revealed that rhizobia primarily influence resistance indirectly via their effect on plant size, raising the possibility that mutualist effects on resistance are common in growth-promoting resource mutualisms. Our results demonstrate that non-defensive mutualists can influence the evolutionary potential of infection-related traits in their hosts. However, if the genotype-by-genotype interactions for infection are common, as we found in this study, the contribution of nutritional mutualisms to heritable variation in infection-related traits will depend on how hosts and mutualists pair up in wild populations.

## Methods

### Study system

*Medicago truncatula* Gaertn. is an annual legume native to the Mediterranean (Cook, 1999). *M. truncatula* plants form association with nitrogen-fixing rhizobia in the soil in the genus *Sinorhizobium*, through the successful exchange of compatible chemical cues and the migration of rhizobia into root tissues. Rhizobia are then housed within nodules that develop from the roots of their host plant. Within these nodules rhizobia fix nitrogen that is traded with the host plant for carbon-containing compounds. The parasitic root knot nematode *Meloidogyne hapla* is an obligate endoparasite with an extremely large host range (Opperman et al., 2008). Female *M. hapla* burrow through intracellular spaces of host roots until they reach a suitable infection site in the vascular cylinder. They then pierce plant cells with their stylet mouthpiece and inject effector proteins that suppress host immune responses and induce cell differentiation producing giant cells from which the nematode siphons nutrients. A gall forms around these infection sites (M. G. K. Jones & Goto, 2011). Previous studies showed that plant maternal families (hereafter, genotypes) and plants inoculated with different rhizobia strains differ in resistance to this parasite (Buxton-Martin et al., 2025; Wood et al., 2018).

We chose 20 accessions of *M. truncatula* from the French National Institute for Agriculture Research (INRAE) germplasm library. We haphazardly chose these accessions to span the range of standing genetic variation in *M. truncatula*, including representative diversity across structured populations as well as resistance to nematode infection and nodulation with rhizobia reported in Wood *et al*. (2018) (see Supplemental for details). We chose 10 strains of *S. meliloti* that were isolated from *M. truncatula* (STable 2). Using FASTANI (Jain et al., 2018), these strains showed a minimum average nucleotide identity of no less than 98% (SFigure 3), indicating all strains belong to a single species. The sources of all *S. meliloti* strains can be found in the Supplementary Materials. We sourced *M. hapla* line LM from the Department of Entomology and Plant Pathology at North Carolina State University, which was originally collected in France (Chen & Roberts, 2003).

### Experimental design and data collection

Following the protocol of Garcia *et al*. (2006), we grew *M. truncatula* in 66 mL Cone-tainer pots (Stuewe & Sons, Inc., Tangent, OR, USA) within a growth chamber at the University of Pennsylvania. Plants were assigned to treatments such that plant genotypes encountered 2-4 rhizobia strains, and each rhizobia strain encountered 5 or 6 host genotypes (Figure 1A). We replicated this experimental design across nematode-infected and nematode-uninfected treatments. Plants were inoculated with nematodes and rhizobia 14 days after planting. Each treatment (plant genotype-rhizobia strain combination) initially included 7-8 replicates, to ensure that at least 4-5 replicates survived to harvesting. See the Supplementary Materials for details regarding inoculation, growth conditions, and sample preparation.

To prevent rhizobia or nematode contamination, each rack of plants contained a single rhizobia treatment and nematode treatment. We distributed rhizobia strain and nematode treatments across different shelves and within the growth chamber in an incomplete block design and randomized plant genotypes within racks. We included shelf location within growth chambers as a fixed-effect blocking variable in our statistical models.

Plants were harvested 52-56 days after planting. Approximately 5-40 nodules were dissected from plant roots during harvesting for a separate experiment and the number of nodules removed was recorded. To harvest, roots were washed in water, and roots and shoots were separated with scissors. Shoots were dried for a minimum of 48 hours in a drying oven at 60 °C and root systems were frozen in -20 °C for later processing.

After drying, shoots were weighed to the nearest tenth of a milligram. To quantify parasitic nematode infection, we dyed the root systems of nematode-infected plants with red food coloring following the methods outlined in Kandouh *et al*. (2019), which stains mature nematode egg sacs. Roots were scanned on an Epson Perfection V850 Pro scanner at 1200 dpi with high contrast background correction following methods outlined in York (2023). Nodules, galls, and nematode egg sacs were manually counted from these images using ImageJ (Schneider et al., 2012). After scanning, roots were dried for >48 hours in a drying oven and weighed to the nearest 0.0001 gram.

### Variance component analyses: Quantifying rhizobia contributions to genetic variation in infection-related traits

To estimate the contribution of rhizobia strains to variation in resistance, virulence, tolerance, and mutualism robustness, we used variance component analyses (Figure 1B). We fit two models for each trait except virulence: one with (Figures 2, 4, 5) and one without (SFigures 5-7) root biomass as a covariate, to capture rhizobia strain effects on infection intensity measured as parasite density (i.e. number of galls per root biomass) and total infection intensity (i.e. number of galls per individual plant). We primarily present models with parasite density but include results that do not correct for root biomass in the Supplementary Materials (SFigures 5-7). Results were qualitatively similar for most analyses.

**Figure 2.**
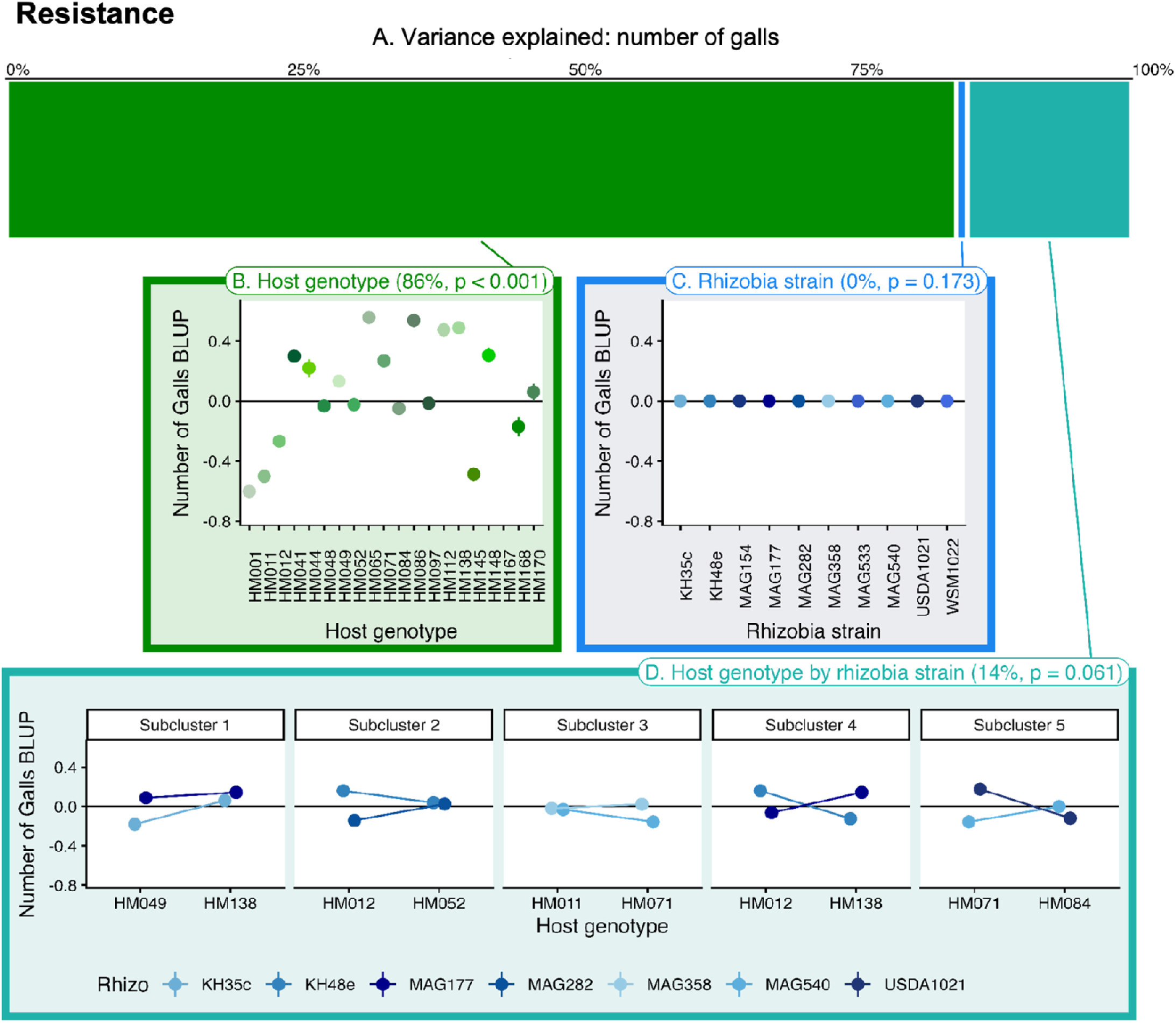
Genetic variation in **resistance** to nematode infection. (A) Variance component analysis of resistance, partitioning variance into three sources: host genotype, rhizobia strain, and the genotype-by-strain interaction. (B-D) Visualizing G_Host_ (B), G_Rhizobia_ (C), and G_Host × Rhizobia_ (D) as variation among host genotypes, rhizobia strains, and a subset of genotype-strain combinations, respectively. Genotype and strain means are the best linear unbiased predictors (BLUPs) for number of galls, extracted from our mixed model (see Methods). Because our experiment was an incomplete factorial design, we only show a subset of plant-rhizobia combinations for the genotype-by-strain interaction (D); see SFigure 2 for subclusters.

To test whether rhizobia strains contributed to variation in resistance to nematode infection, we fit mixed-effects models with number of galls as the response variable and rhizobia strain, host genotype, and the genotype-by-strain interaction as random effects. This analysis only included nematode-infected plants. In this analysis, a significant effect of rhizobia strain or significant genotype-by-strain interaction indicates that rhizobia contribute to genetic variation in resistance via indirect genetic effects (G_Rhizobia_) or intergenomic epistasis (G_Host × Rhizobia_) respectively. Experimental block was included as a fixed effect, and we fit a negative binomial error distribution (Lindén & Mäntyniemi, 2011).

Virulence is measured as the difference in fitness between infected and uninfected hosts. To test whether rhizobia strains contributed to variation in virulence, we fit a mixed-effect model with shoot biomass, our host fitness proxy, as the response variable and rhizobia strain, host genotype, genotype-by-strain, and all three of these effects interacting with nematode infection status as random effects. Experimental block and nematode infection status were included as fixed effects. In this analysis, a significant effect of infection-by-strain indicates rhizobia strain effects on virulence and a significant effect of infection-by-strain-by-genotype indicates that rhizobia contribute to virulence via indirect genetic effects (G_Rhizobia_) or intergenomic epistasis (G_Host × Rhizobia_) respectively.

Tolerance is the relationship between infection intensity and host fitness. To test whether rhizobia strains contributed to variation in tolerance, we fit mixed-effects models with shoot biomass as the response variable and rhizobia strain, host genotype, and both of these effects interacting with the number of galls as random effects. Unlike our other analyses, because each genotype-by-strain combination had 4-8 replicates, we did not have the statistical power to confidently estimate slopes for specific host genotype-rhizobia strain combinations and therefore did not include genotype-by-strain or infection-by-genotype-by-strain effects in our model. For random effects interacting with the number of galls, we included random slopes and no random intercept for the number of galls. We included the number of galls and experimental block as fixed effects. In this analysis, a significant interaction between number of galls and rhizobia strain indicates that rhizobia strain contributes to genetic variation in tolerance via indirect genetic effects (G_Rhizobia_).

We defined mutualism robustness as the difference in nodulation between infected and uninfected hosts. To test whether rhizobia strains contributed to variation in mutualism robustness, we fit mixed-effects models with the number of nodules as the response variable and rhizobia strain, host genotype, genotype-by-strain, and all three of these effects interacting with nematode infection status as random effects. Experimental block and nematode infection status were included as fixed effects. In this analysis, a significant effect of infection-by-strain or infection-by-strain-by-genotype indicates rhizobia strain effects on mutualism robustness via indirect genetic effects (G_Rhizobia_) or intergenomic epistasis (G_Host × Rhizobia_) respectively. We used a negative binomial distribution error (Lindén & Mäntyniemi, 2011).

We implemented our data analysis in R. We fit mixed-effects models using the glmmTMB package (Brooks et al., 2017), and used the Sattherthwaite’s degrees of freedom method implemented in the lmerTest package (Kuznetsova et al., 2017) to estimate p-values and infer statistical significance. We used the DHARMa package (Florian, 2016) to check that model residuals were normally distributed and homoscedastic. We tested significance of random effects with likelihood ratio tests. We used the MuMIn package (Bartoń, 2025) to calculate the conditional and marginal R^2^ values for each model and used the trigamma method when possible per the package author’s suggestion.

### Path analysis: Distinguishing direct and indirect effects of rhizobia on infection intensity

Resource mutualists like rhizobia could affect the intensity of parasite infection (i.e., the number of parasites on a host) directly, or indirectly via their effects on host growth, which impacts the amount of host tissue available to infect. We therefore used path analysis to ask whether the effect of rhizobia on resistance was mediated by root biomass. Our path analysis used the following models:

### Outcome model: Number of galls ∼ Root biomass + Host genotype + Rhizobia strain Mediating model: Root biomass ∼ Host genotype + Rhizobia strain

To test for significant effects in the overall path analysis, we used a negative binomial GLM implemented with the glm.nb function from the MASS package in R (Venables & Ripley, 2002) to fit our outcome models that had gall number as the response variable (Lindén & Mäntyniemi, 2011) and a Gaussian linear model implemented in the lm function from the STAT package (Bolar, 2019) to fit the mediating model. To estimate standardized path coefficients we used a Gaussian distribution for both models fit with the lm function. We used the piecewiseSEM package (Lefcheck et al., 2025) to test and structure our path analysis. We used sum-to-zero contrasts in all our models. To test for the significance of direct and indirect effects in the per strain path analyses, we used the boot function in the boot package (Canty & Ripley, 2025) to perform bootstrapping analysis with 10,000 resamples to acquire confidence intervals for the product of path coefficients and the proportion of meditation using the Gaussian models.

## Results

### Resistance

Our resistance model (Figure 2) explained 41.4% of the total variation in the number of galls (R^2^_C_). Of the total variation in the number of galls, 10.28% was explained by fixed effects (R^2^_M_) leaving 31.12% explained by our random effects. Of the variation in the number of galls explained by random effects, our variance component analysis attributed 86% to host genotype (p < 0.0001), 14% to the genotype-by-strain interaction (p = 0.0610), and no significant variation to rhizobia strain (p = 0.1730). These results indicate that host genotype was the largest determinant of resistance but that genotype-by-strain interaction does contribute a significant variation to resistance. Our variance component analysis that did not correct for root biomass indicated that only host genotype contributed to variation in gall numbers.

### Virulence

Our virulence model (Figure 3) explained 92.6% of the total variance in shoot biomass (host fitness) (R^2^_C_). Of the total variance in shoot biomass, 27.51% was explained by fixed effects (R^2^_M_) leaving approximately 65.09% explained by our random effects. Of the variation in shoot biomass explained by random effects, 5% was attributed to the infection-by-strain interaction (p = 0.048). The infection-by-genotype interaction and infection-by-genotype-by-rhizoba-strain interaction explained 3% and 8% of the variance, respectively, but neither was statistically significant (p = 0.133 and p = 0.222 respectively). These results indicate that rhizobia strains contribute to variation in the fitness consequences of nematode infection, but that their effect depends on the genotype of their host plant.

**Figure 3.**
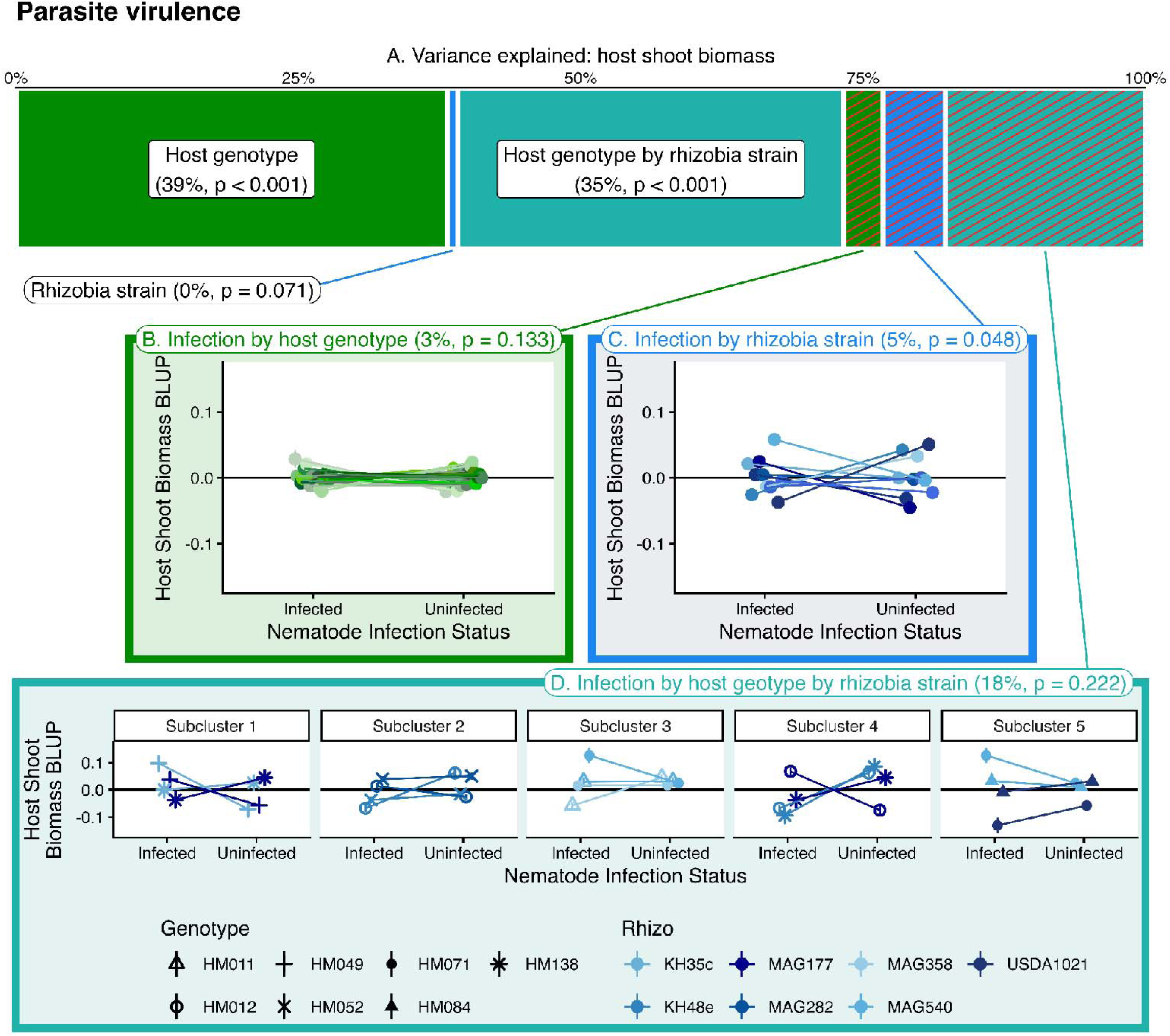
Genetic variation in **parasite virulence**. (A) Variance component analysis of virulence, partitioning variance into three sources: host genotype (the infection x genotype interaction), rhizobia strain (the infection x strain interaction), and the genotype-by-strain interaction (the three-way interaction). (B-D) Visualizing G_Host_ (B), G_Rhizobia_ (C), and G_Host × Rhizobia_ (D) as variation among host genotypes, rhizobia strains, and genotype-strain combinations, respectively, in the fitness cost of infection. Genotype and strain means are the best linear unbiased predictors (BLUPs) extracted from our mixed model (see Methods). Because our experiment was an incomplete factorial design, we only show a subset of plant-rhizobia combinations for the genotype-by-strain interaction (D); see SFigure 2 for subclusters.

### Tolerance

Our tolerance model that corrected for root biomass (Figure 4) explained 77.18% of the total variation in shoot biomass (host fitness) (R^2^_C_). Of the total variation in shoot biomass, 69.57% was explained by fixed effects (R^2^_M_) leaving approximately 7.61% explained by our random effects. Of the variation in shoot biomass explained by random effects, our variance component analysis attributed no variance to effects interacting with nematode infection severity (number of galls). Instead, all variation in host fitness was attributed to host genotype (100%, p < 0.0001). Qualitatively similar results were found from our tolerance model that did not correct for root biomass (R^2^_C_= 0.2253; R^2^_M_= 0.0697). These results indicate that rhizobia strains do not contribute to variation in tolerance.

**Figure 4.**
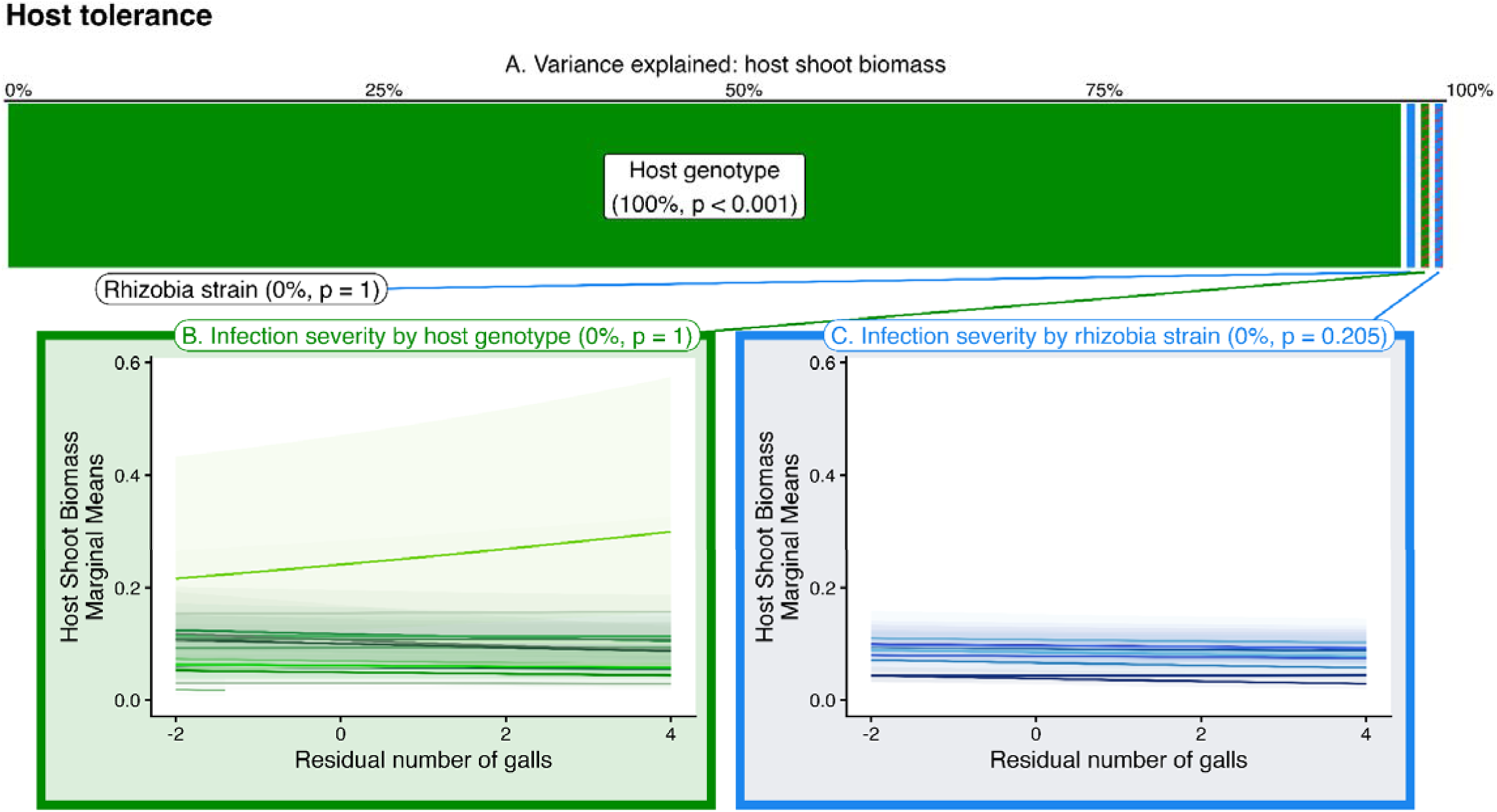
Genetic variation in **tolerance**. (A) Variance component analysis of virulence, partitioning variance in tolerance into three sources: host genotype (the galls x genotype interaction) and rhizobia strain (galls x strain interaction). We did not test for a genotype-by-strain interaction for tolerance. (B-C) Visualizing G_Host_ (B) and G_Rhizobia_ (C) as variation among host genotypes and rhizobia strains, respectively, in the relationship between parasite load and fitness. No significant infection-by-strain interactions were observed, indicating no effect of rhizobia strain on host tolerance.

### Mutualist robustness

Our mutualism robustness model that corrected for root biomass (Figure 5) explained 46.86% of the total variation in nodule counts (R^2^_M_). Of the total variation in nodule counts, 11.71% was explained by fixed effects (R^2^_C_) leaving approximately 35.15% explained by random effects. Of the variation in nodule counts explained by random effects, 12% was attributed to the infection-by-genotype interaction (p = < 0.0001) and 16% to the infection-by-genotype-by-strain interaction (p = 0.0480). No variance was attributed to the infection-by-strain interaction (p = 0.5409). Our mutualism robustness model that did not correct for root biomass showed qualitatively similar results. These results indicate that individual host genotype and rhizobia strain combinations contribute to variation in mutualism robustness to parasite infection, but that the rhizobia effect depends on the genotype of its host.

**Figure 5.**
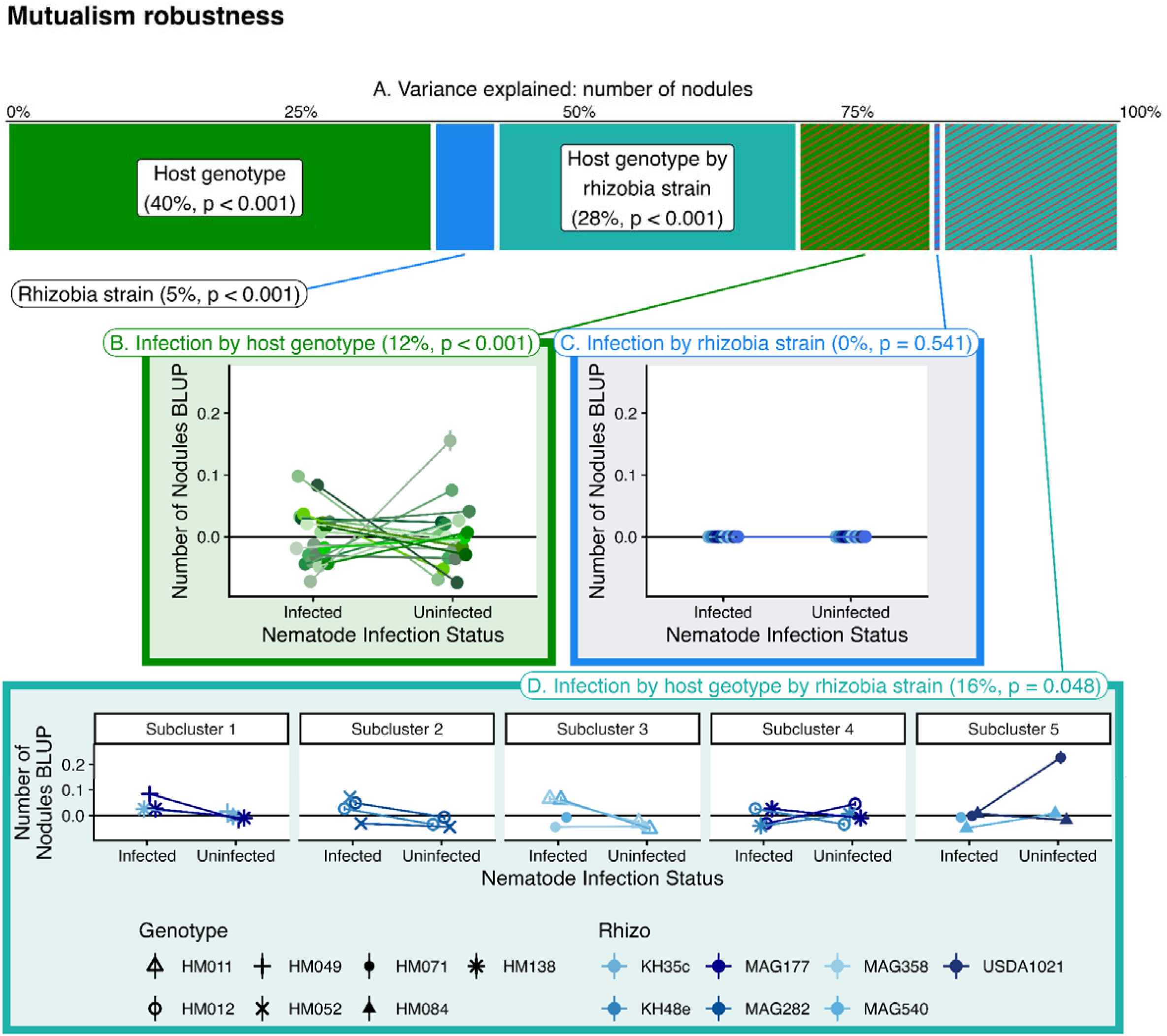
Genetic variation in **mutualism robustness to infection**. (A) Variance component analysis of virulence, partitioning variance into three sources: host genotype (the infection x genotype interaction), rhizobia strain (the infection x strain interaction), and the genotype-by-strain interaction (the three-way interaction). (B-D) Visualizing G_Host_ (B), G_Rhizobia_ (C), and G_Host × Rhizobia_ (D) as variation among host genotypes, rhizobia strains, and genotype-strain combinations, respectively, in the impact of parasite infection on nodulation. Genotype and strain means are the best linear unbiased predictors (BLUPs) extracted from our mixed model (see Methods). Because our experiment was an incomplete factorial design, we only show a subset of plant-rhizobia combinations for the genotype-by-strain interaction (D); see SFigure 5 for subclusters.

### Rhizobia effects on nematode infection intensity are primarily mediated by their effect on plant size

In our path analysis (Figure 6), we found that rhizobia strains varied in their influence on root biomass (p<0.0001) but not on gall numbers (p=0.1381) (Figure 2). Gall numbers increased with root biomass (β=0.3151, p<0.0001). These results indicate that after correcting for the differences in host genotype, larger roots are associated with greater number of galls, and rhizobia differ in their effect on root growth. These path analysis results are consistent with the difference between variance component analyses that did (Figure 2) and did not (SFigure 5) correct for root biomass and indicate that their effect on plant growth is one mechanism through which rhizobia strains influence resistance to nematodes.

**Figure 6:**
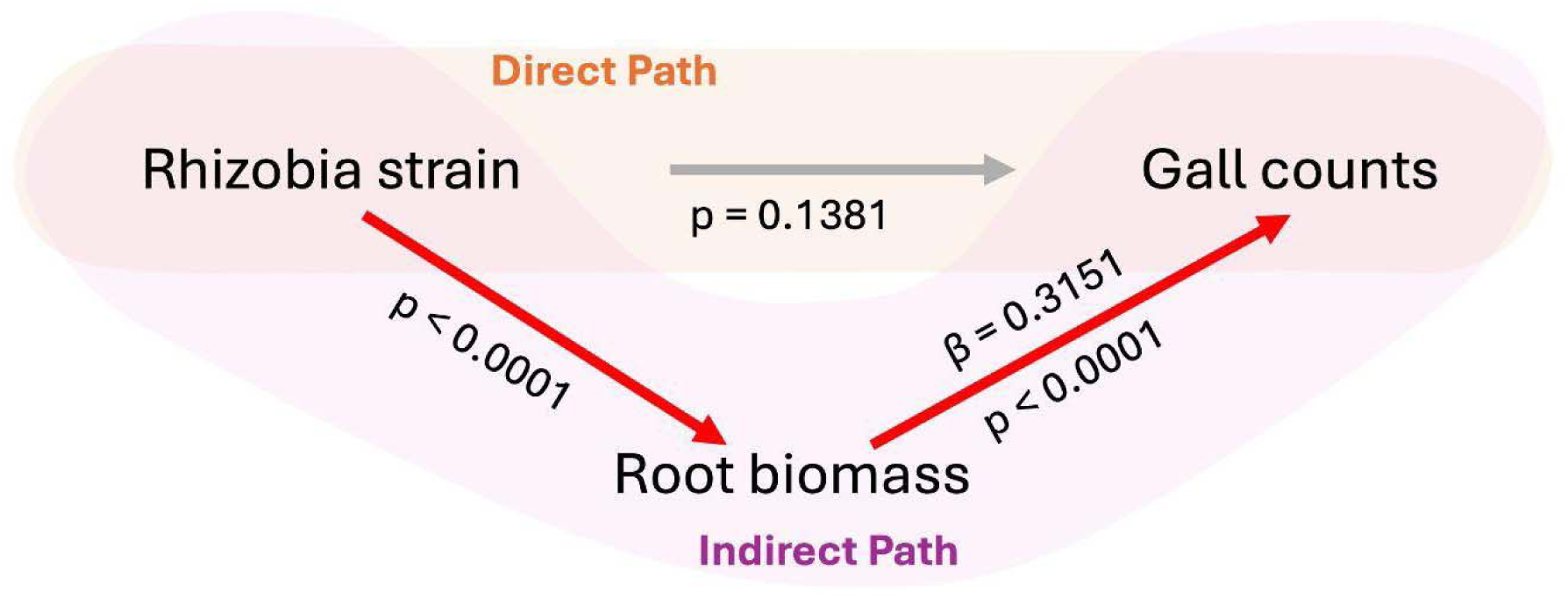
Path analysis results showing significant effect of rhizobia strain on root biomass (p < 0.0001) and a significant effect of root biomass on the number of galls (β=0.3151, p < 0.0001) but no significant direct effect of rhizobia strains on the number of galls (p = 0.1381). This path analysis corrected for variation in these traits among host genotypes (see Methods).

## Discussion

Here we report that nitrogen-fixing rhizobia contribute to genetic variation in infection-related traits in their host plant. These effects manifested as differences in parasite virulence among plants inoculated with different rhizobia strains (i.e., G_Rhizobia_), as well as intergenomic epistasis for resistance (i.e., G_Host × Rhizobia_) (Table 1). To our surprise, we detected no significant host genetic variance for parasite virulence, only rhizobia strain and host-genotype-by-rhizobia-strain effects.The rhizobia effect on resistance is partially explained by a core property of this nutritional mutualist: its effect on plant growth. In addition to these canonical infection-related traits, genotype-by-genotype interactions between plants and rhizobia also contributed to genetic variation in mutualism robustness to parasite infection. These results add to the growing literature on the ecological and evolutionary significance of symbiotic extended phenotypes, and suggests that non-defensive mutualists may be overlooked contributors to the evolutionary potential of defense traits in their hosts.

**Table 1.**
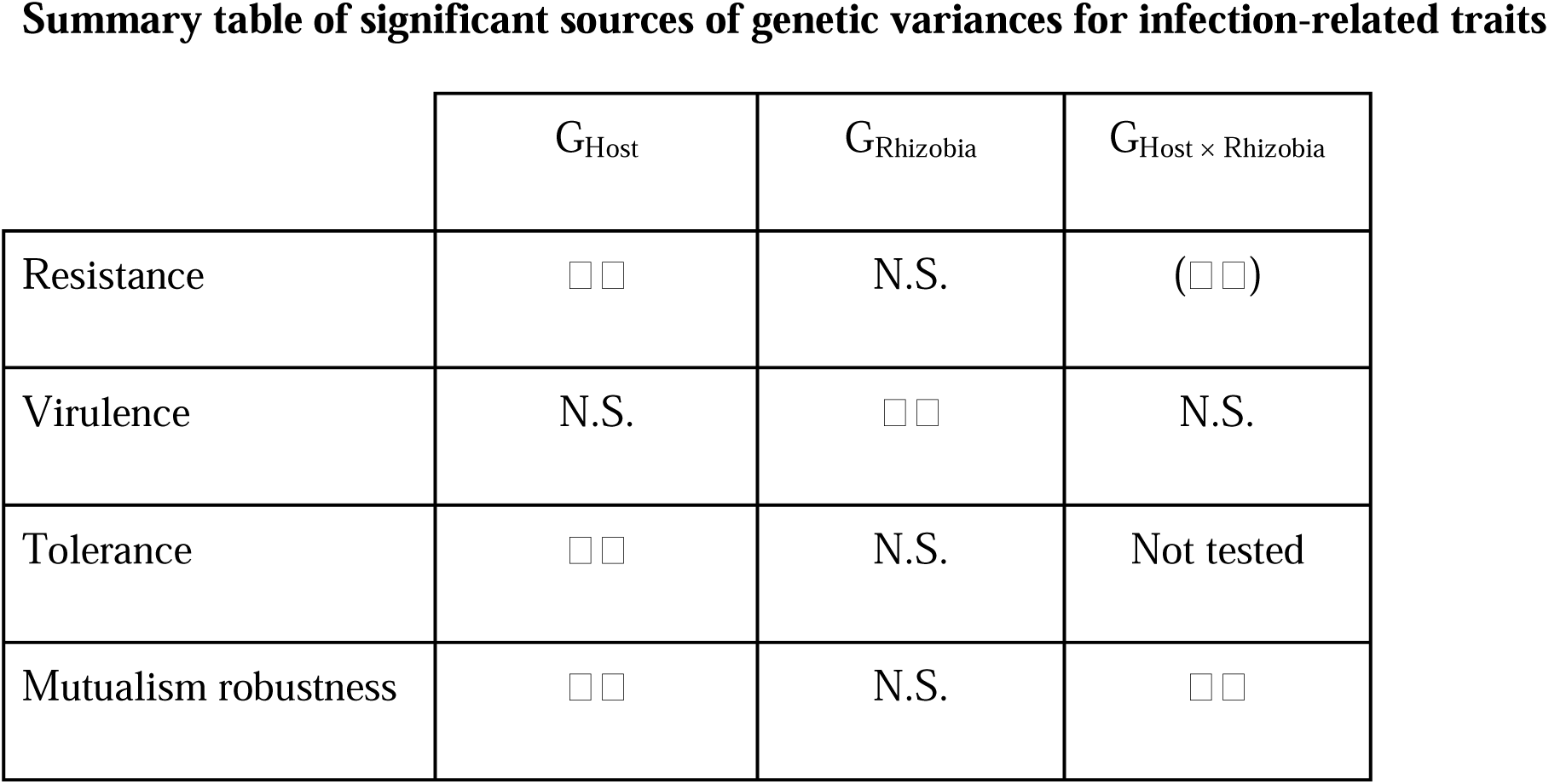
A summative table showing whether genetic variances attributed to host genotype, rhizobia strain, or genotype-by-strain were found for each infection-related traits tested. Check marks indicate a significant effect (p<0.05), check marks within parentheses indicate a marginally significant (p<0.1) effect, and N.S. indicates no significant effect (p>0.1).

### Rhizobia contributed to genetic variation in resistance and virulence

Rhizobia contributed significantly to genetic variation in two of the three canonical infection-related traits in our study: resistance (total parasite load) and virulence (the difference in fitness between infected and uninfected hosts). The contribution of rhizobia to genetic variation in resistance was small relative to the host, and completely tied up in intergenomic epistasis (G_Host × Rhizobia_). Overall, variation in resistance was primarily governed by the host genome (G_Host_ = 86% of total genetic variance; Figure 2). By contrast, rhizobia contributed substantially to genetic variation in virulence. Our variance component analyses attributed 5% of variation in virulence to rhizobia (the nematode infection-by-rhizobia strain interaction for plant biomass) and an additional 18% of variation to the genotype-by-genotype interaction (GxG) between plants and rhizobia (the infection-by-genotype-by-strain interaction for plant biomass) (although this term was not statistically significant; Figure 3). To our surprise, only 3% of variation in virulence was attributed to the host alone (infection-by-plant genotype interaction for plant biomass). In other words, rhizobia contributed to nearly 90% of total genetic variation in virulence. These results show that rhizobial strain identity is a primary determinant of the virulence of nematode infections in *M. truncatula*.

Significant genotype-by-genotype interactions (GxG) for resistance are consistent with the established prevalence of intergenomic epistasis in both nutritional and defensive mutualisms (Batstone et al., 2022; Heath, 2010; Heath & Tiffin, 2007; Parker et al., 2017; Schmid et al., 2012). Intergenomic epistasis is hypothesized to contribute to the maintenance of genetic variation in mutualisms, because it creates a situation where the mutualist strain favored by selection differs across host genetic backgrounds (and vice versa) (Heath & Stinchcombe, 2014). Ultimately, though, the evolutionary ramifications of GxG depend on how hosts and symbionts partner in wild populations (De Lisle et al., 2022). If plants and rhizobia pair non-randomly, intergenomic epistasis could contribute to additive genetic variance in parasite resistance, because a rhizobia strain’s effect on resistance will only occur in a subset of host genetic backgrounds. Non-random pairing between host and symbiont could arise from partner choice, which is common in the legume-rhizobia mutualism (Simms & Taylor, 2002). However, to the best of our knowledge, the covariance between host and symbiont genotypes has not been directly measured in a facultative mutualism (De Lisle et al., 2022).

Finally, we observed no rhizobia strain effects on tolerance. Tolerance and resistance are two strategies that hosts evolve in response to parasites, each with differing evolutionary consequences. Increased resistance tends to select for increased virulence, potentially leading to an evolutionary arms race whereas tolerance does not (Vorburger & Perlman, 2018). Examples of symbionts that protect their hosts via increasing tolerance appear to be rare or underrepresented in the literature (Vorburger & Perlman, 2018) and so the lack of rhizobial effects on host tolerance may support this notion that microbial effects on host tolerance may be rare. However, resource availability can influence tolerance (Budischak & Cressler, 2018; Yi et al., 2025), making this lack of rhizobial effect on tolerance somewhat surprising.

### Rhizobia contribute to genetic variation in mutualism robustness to parasite infection

Previous research found that, in addition to stealing resources from their host, parasitic nematodes decrease nodulation with rhizobia. The magnitude of this effect varied across plant genetic backgrounds, indicating that mutualism robustness is a heritable plant trait (Wood et al., 2018). In the present study, we replicated this result, finding significant host genetic variation for mutualism robustness (infection-by-host genotype interaction for nodulation; Figure 5).

Furthermore, here we found that mutualism robustness is also influenced by a genotype-by-genotype interaction between plants and rhizobia (infection-by-genotype-by-strain interaction for nodulation; Figure 5). Rhizobia thus contain genetic variation for mutualism robustness in the face of nematode infection, indicating that rhizobia evolution may influence mutualism robustness. This result is consistent with the established importance of context dependence in mutualisms (Carlson et al., 2025; Chamberlain et al., 2014; Heath & Tiffin, 2007; Parker et al., 2017). It suggests that parasite infection may be an important environmental factor shaping mutualism expression and evolution, and suggests that the mutualism disruption of mutualisms may be an overlooked cost of infection

### The genetic and physiological basis of strain effects on parasite infection

Mutualist effects on parasite infection could arise from two underlying mechanisms: immune crosstalk between mutualism and parasitism, or resource competition or facilitation (Carlson et al., 2025). While our experiment cannot directly discriminate between immune- and resource-mediated mechanisms, our path analysis (Figure 6) provides a basis for hypothesizing about the biological processes that may mediate the effects we observed. We used this analysis to evaluate whether the rhizobia effect on nematode resistance is mediated by the rhizobia effect on plant growth (root biomass), consistent with a resource-mediated link between rhizobia and parasite resistance. Our results are consistent with this hypothesis: rhizobia differed in their effect on root biomass, and root biomass was associated with parasite load, suggesting that the rhizobia effect on parasite load is associated with the resources they provide. This suggests that strain effects on parasite resistance may arise from genotype-by-genotype interactions between plants and rhizobia for resource- or growth-related biological processes.

By contrast, the direct path linking rhizobia and parasite load was not significant. One possible interpretation of this result is that immune crosstalk between rhizobia and nematodes did not substantially influence nematode resistance in this study. However, in our variance component analyses that corrected for plant size (i.e., analyses that included root biomass as a covariate), we still detected a significant rhizobia contribution to genetic variation in resistance, via GxG between rhizobia and host. Together, these observations suggest that, at least for some strains on some plant genetic backgrounds, the rhizobia effect on parasite resistance is not fully explained by the resources they provide. This inference is also supported by our recent transcriptomic study showing that rhizobia strains promote differential expression of genes enriched for immune functions in nematode galls (Buxton-Martin et al., 2025).

Ultimately, deciphering the biological basis involved in rhizobial strain effects on infection-related traits is important because these mechanisms differ in their implications for host-parasite evolution. In defensive mutualisms, the mechanisms by which defensive symbionts affect virulence (Armitage et al., 2022; Vorburger & Perlman, 2018) and the costs associated with harboring defensive mutualists (King & Bonsall, 2017) influence the evolutionary consequences of those protective effects. Competition between parasites and mutualists for resources is expected to in most cases increase virulence; interference competition is expected to decrease virulence, and effects mediated by immune responses may vary significantly in their evolutionary consequences for virulence (Armitage et al., 2022; Vorburger & Perlman, 2018).

Our path analysis also highlights an important consideration for researchers studying interactions between parasites and resource mutualists: is absolute parasite load or parasite density (i.e., body size-adjusted parasite load) a more appropriate metric of infection severity? While absolute parasite load may provide ecologically relevant information about parasite populations, parasite density may be a better indicator of the fitness cost of parasite infection. If larger hosts typically harbor more parasites, resource mutualists may often increase absolute parasite load because they facilitate host growth. However, their effects on parasite density and the net fitness cost of infection may be less predictable. With this in mind we recommend considering carefully how infection-related traits – especially infection intensity – is quantified when in systems in which resource mutualists impact host growth.

### Ecological and evolutionary implications of nutritional mutualist effects on heritable variation in infection-related traits

Our study shows that mutualist contributions to genetic variation in infection-related traits are not restricted to defensive mutualisms, i.e., interactions where the primary benefit is defense. This is consistent with pervasive effects of microbial symbionts on a wide range of host traits (Friesen et al., 2011). Studies in defensive mutualisms suggest that these effects are evolutionarily significant: defensive mutualists contribute substantially to heritable variation in infection related traits (Martinez et al., 2017; Parker et al., 2017) and they can drive the rapid evolution and coevolution of resistance, tolerance, and virulence (Ford et al., 2016, 2017; May & Nelson, 2014; Rafaluk-Mohr et al., 2018, 2022; Vorburger & Perlman, 2018). Our study adds to the growing body of studies showing that these effects may not be restricted to defensive mutualisms (Wood et al., *in review*). Nitrogen-fixing rhizobia are textbook nutritional mutualists. Yet our study and several others show that they can affect their host’s interactions with enemies (Irmer et al., 2015; Prévitali et al., 2025; Simonsen & Stinchcombe, 2014; Wood et al., 2018).

Future studies should directly test whether genetic variation in non-defensive mutualists contributes to the evolution of defense traits in their hosts, as it does in defensive mutualisms. Non-defensive mutualists could accelerate defense evolution by increasing genetic variation available to selection in hosts, or directly via the rapid evolution of microbial extended phenotypes (De Lisle et al., 2022; Henry et al., 2021; O’Brien et al., 2021; Wood et al., *in review*). However, if the underlying microbial genes are subject to conflicting selection (for example, divergent selection between the free-living and symbiotic life stages), then joint trait evolution will be difficult to predict without understanding how selection operates in both the host and the microbe. Therefore, understanding how selection acts on symbiotic extended phenotypes could be key to predicting the evolution of joint traits, including those related to parasite infection. Furthermore, pervasive GxG between hosts and mutualists means that total genetic variation in a joint trait like defense depends on how hosts and symbionts partner in wild populations. Rhizobia could actually *decrease* total genetic variance in the plants response to infection if GxG masks heritable differences among host genotypes (Heath, 2010; Wood et al., *in review*).

Finally, our study raises several additional questions. First, are non-defensive mutualists more likely to influence resistance against specific types of antagonists? Understanding the immune- and resource-mediated mechanisms underlying nutritional mutualist effects on defense traits might allow us to predict when mutualists will affect interactions with antagonists, and if so, which ones. Microbial mutualists may be especially likely to influence defense against antagonists governed by similar immune pathways, or defenses whose cost overlaps with the resources provided or consumed by the mutualist. For example, in our system, nematodes and rhizobia have significant overlapping pathways by which they interact with their host (Koltai et al., 2001). Finally, our results highlighted the importance of intergenomic epistasis between hosts and symbiotic mutualists, but we did not consider GxG between the symbionts themselves. Our experiment involved single-strain inoculations, yet plants are can be colonized by multiple rhizobia strains and the benefit that the rhizobia provide depends on the strain(s) they are co-inoculated with (Rahman et al., 2023; Westhoek et al., 2021). This suggests that other symbiotic extended phenotypes, including rhizobia effects on infection-related traits, may also be influenced by GxG between rhizobia strains. These lines of inquiry parallel the emerging focus within parasitology and phytopathology on concepts like the pathobiome (Vayssier-Taussat et al., 2014) and disease tetrahedron (Leveau, 2024), which incorporate ecological realism into host-parasite interactions.

