## Supplemental Information for "Mutualistic rhizobia harbor genetic variation for traits related to parasite infection"

**Methods S1**: Sample preparation

Per Garcia *et al*. (2006), we mechanically scarified *M. truncatula* seeds with sandpaper, surface sterilized them with 10% bleach solution, stratified seeds by putting them in the dark at 4 °C for 2 days in petri dishes with paper disks saturated with sterile water, and then incubated them at room temperature for 2 days. We planted germinated seedlings in 120 mL Cone-tainers filled with 1:4 mixture of perlite and sand and autoclaved twice with >24 hours between each autoclaving (Barker et al., 2006). We maintained seedlings in growth chambers at the University of Pennsylvania. Growth chambers were kept in a 16:8 light:dark cycle, alternating between 25 °C during the light cycle and 21 °C during the dark cycle. After planting we fertilized plants on a recurring basis (details of the fertilizing procedure and schedule found in SI Methods).

We used three different fertilizers containing 5 mM N, 0.625 mM N, or 0 mM N. Recipes for fertilizing are found in SI Table S1 and are modified from the recipes found in Batstone *et al.* (2017) and Moreau *et al.* (2008). Plants were fertilized three times a week. On the day of planting, we fertilized all plants with 5 mL of 5 mM fertilizer and 10 mL of diH2O. The three subsequent days of fertilizing we used 5 mL of 0.625 mM N fertilizer and 10 mL of diH2O. For every following day of fertilizing we used 0 mM N fertilizer and 10 mL of diH2O.

On day 14 after planting, we inoculated each M. truncatula plant with rhizobia and nematode treatments. Nematode inoculation consisted of 5 mL of double deionized water solution containing 86 nematode eggs mL-1 (~430 eggs total). Rhizobial inoculation consisted of 5 mL of 2 day old liquid culture in tryptone yeast media, diluted to OD600 of 0.1 (Journet et al., 2006). Plants not inoculated with nematodes were given 5 mL of double deionized water. Plants were fertilized on subsequent days with details of the fertilizing procedure and schedule found in SI Methods.

**Methods S2**: Population structure analysis

To ensure that the host genotypes are representative of the range of standing genetic variation in *M. truncatula*, we looked for structured populations within the Medicago HapMap. We took the SNPs from the HapMap for various host genotypes, pruned them with PLINK (noa) removing linked and intragenic SNPs, and then performed a principal component analysis with the resulting SNPs. Three distinct subclusters were apparent, indicating population structure within the HapMap. We overlayed rate of nodulation and rate of nematode infection data according to Wood et al. (2018). The resulting PCA (SFigure 1) was used to pseudorandomly select across population structure, willingness to partner with rhizobia, and susceptibility to nematodes.

**Table S1**: Table showing the constituents of the various fertilizer solutions used in this experiment.

| Solution | Molarity  (mol / L) | 1 L of 0 mM N Fertilizer (mL) | 1 L of 0.625 mM N Fertilizer (mL) | 1 L of 5 mM N Fertilizer (mL) |
| --- | --- | --- | --- | --- |
| MgSO_4_ * | 0.5 | 4 | 4 | 4 |
| CaCl_2_ * | 1 | 5 | 4.530 | 3.125 |
| Fe-EDTA | 0.02 | 2.5 | 2.5 | 2.5 |
| MnSO_4_ | 0.006 | 0.1 | 0.1 | 0.1 |
| CuSO_4_ | 0.006 | 0.1 | 0.1 | 0.1 |
| ZnSO_4_ | 0.006 | 0.1 | 0.1 | 0.1 |
| H_3_BO_3_ | 0.016 | 0.1 | 0.1 | 0.1 |
| Na_2_MoO_4_ | 0.005 | 0.1 | 0.1 | 0.1 |
| KNO_3_ | 0.625 | 0 | 0.25 | 1 |
| Ca(NO_3_)_2_ | 0.938 | 0 | 0.5 | 2 |
| NaNO_3_ | 2.5 | 0 | 0 | 1 |
| K_2_HPO_4_ | 1.2 | 2 | 2 | 2 |
| K_2_SO_4_ * | 0.43 | 2.197 | 1.953 | 1.802 |
| NaCl | 0.2 | 1 | 1 | 1 |
| diH_2_O | - | 982.803 | 982.767 | 981.073 |

**Table S2**: Table showing rhizobia strain origination and other information.

| **Strain ID** | **NCBI Genome Assembly** | **NCBI Genome Submitter** | **Date** | **Associated Citation** |
| --- | --- | --- | --- | --- |
| USDA1021 | ASM219744v1 | University of Minnesota | June 16, 2017 | Sagawara *et al.* 2013 |
| WSM1022 | ASM1331577v1 | University of Warwick | June 10, 2020 | Terpolilli *et al.* 2008 |
| MAG358 | ASM3371665v1 | University of Minnesota | Nov 15, 2023 | Riley *et al.* 2023 |
| MAG282 | ASM3371661v1 | University of Minnesota | Nov 15, 2023 | Riley *et al.* 2023 |
| MAG154 | ASM3371645v1 | University of Minnesota | Nov 15, 2023 | Riley *et al.* 2023 |
| MAG533 | ASM3371425v1 | University of Minnesota | Nov 15, 2023 | Riley *et al.* 2023 |
| MAG540 | ASM3371612v1 | University of Minnesota | Nov 15, 2023 | Riley *et al.* 2023 |
| MAG177 | ASM3371563v1 | University of Minnesota | Nov 15, 2023 | Riley *et al.* 2023 |
| KH35c | ASM219710v1 | University of Minnesota | June 16, 2017 | Sugawara *et al.* 2013 |
| KH48e | ASM959994v1 | Medicago HapMap Project | Nov 5, 2019 | Sugawara *et al.* 2013 |

Riley, A. B., Grillo, M. A., Epstein, B., Tiffin, P., & Heath, K. D. (2023). Discordant population structure among rhizobium divided genomes and their legume hosts. Molecular ecology, 32(10), 2646-2659.

Sugawara, M., Epstein, B., Badgley, B. D., Unno, T., Xu, L., Reese, J., ... & Sadowsky, M. J. (2013). Comparative genomics of the core and accessory genomes of 48 Sinorhizobium strains comprising five genospecies. Genome biology, 14(2), R17.

Terpolilli, J. J., O’Hara, G. W., Tiwari, R. P., Dilworth, M. J., & Howieson, J. G. (2008). The model legume Medicago truncatula A17 is poorly matched for N2 fixation with the sequenced microsymbiont Sinorhizobium meliloti 1021. New Phytologist, 179(1), 62-66.

**Figure S1:** Population structure in host genotype
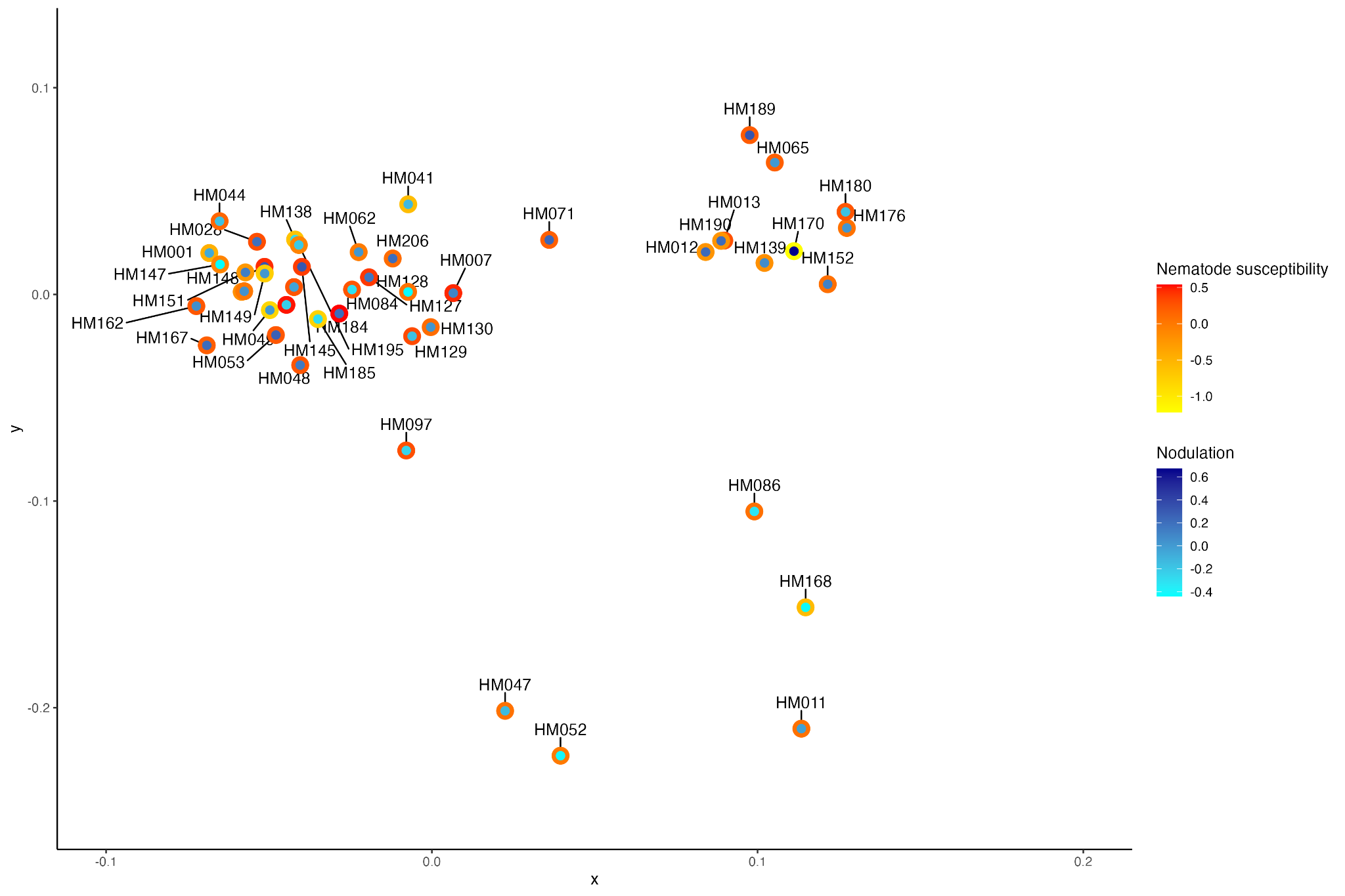

**Figure S2:** Subclusters in the experimental design from which genotype-by-strain effects can be visualized

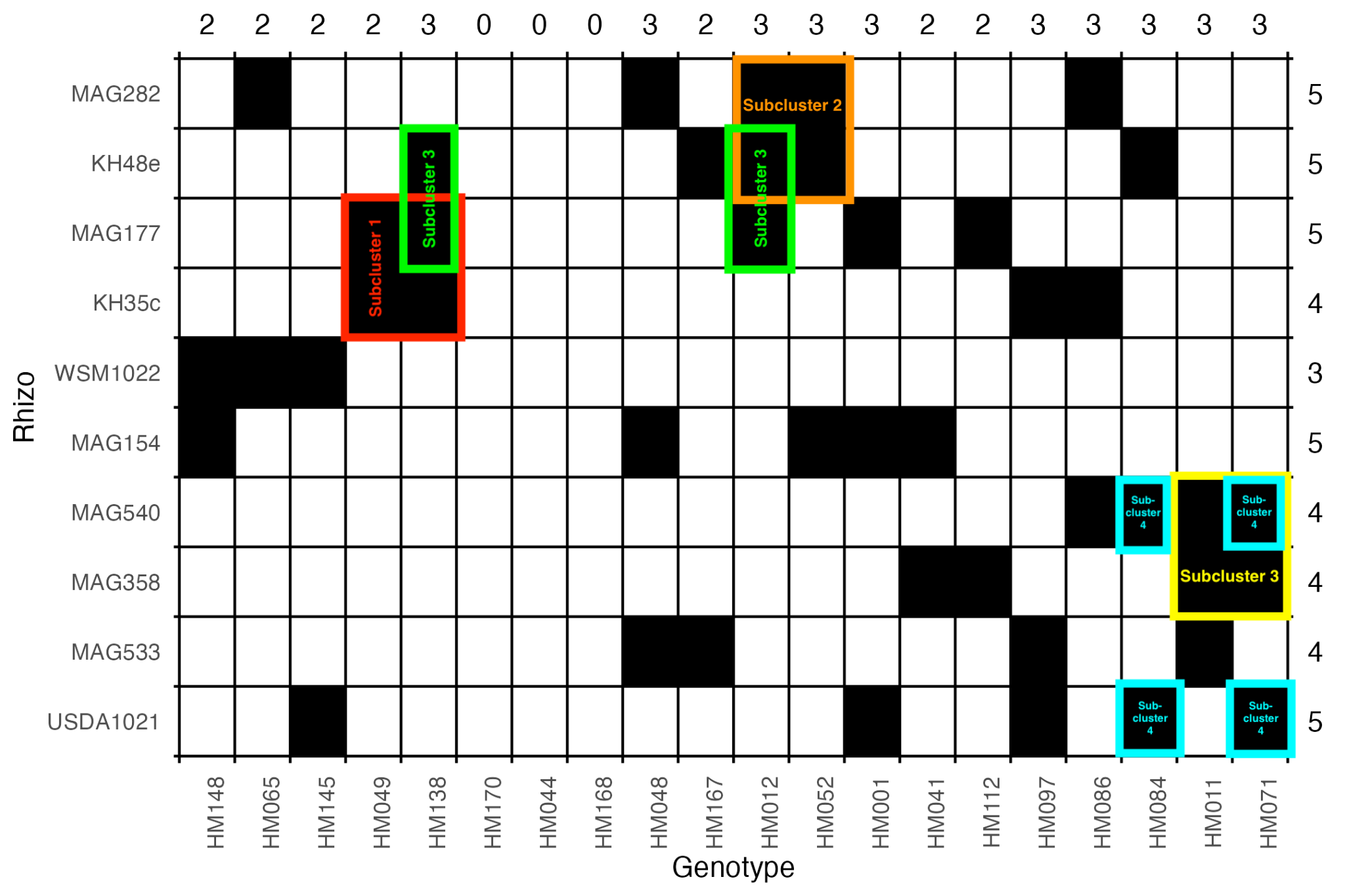

**Figure S3:** FASTANI relatedness between rhizobia strains

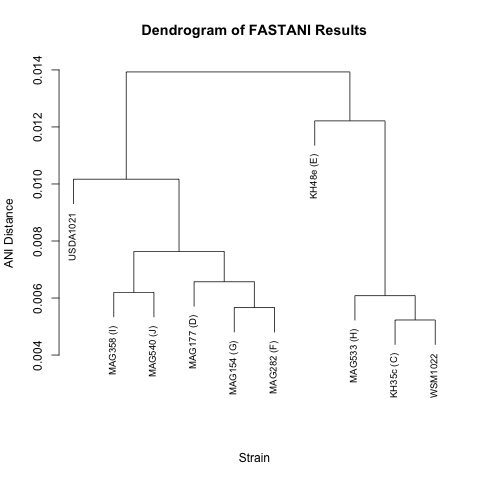

**Figure S4:** Correlation between host trait data.

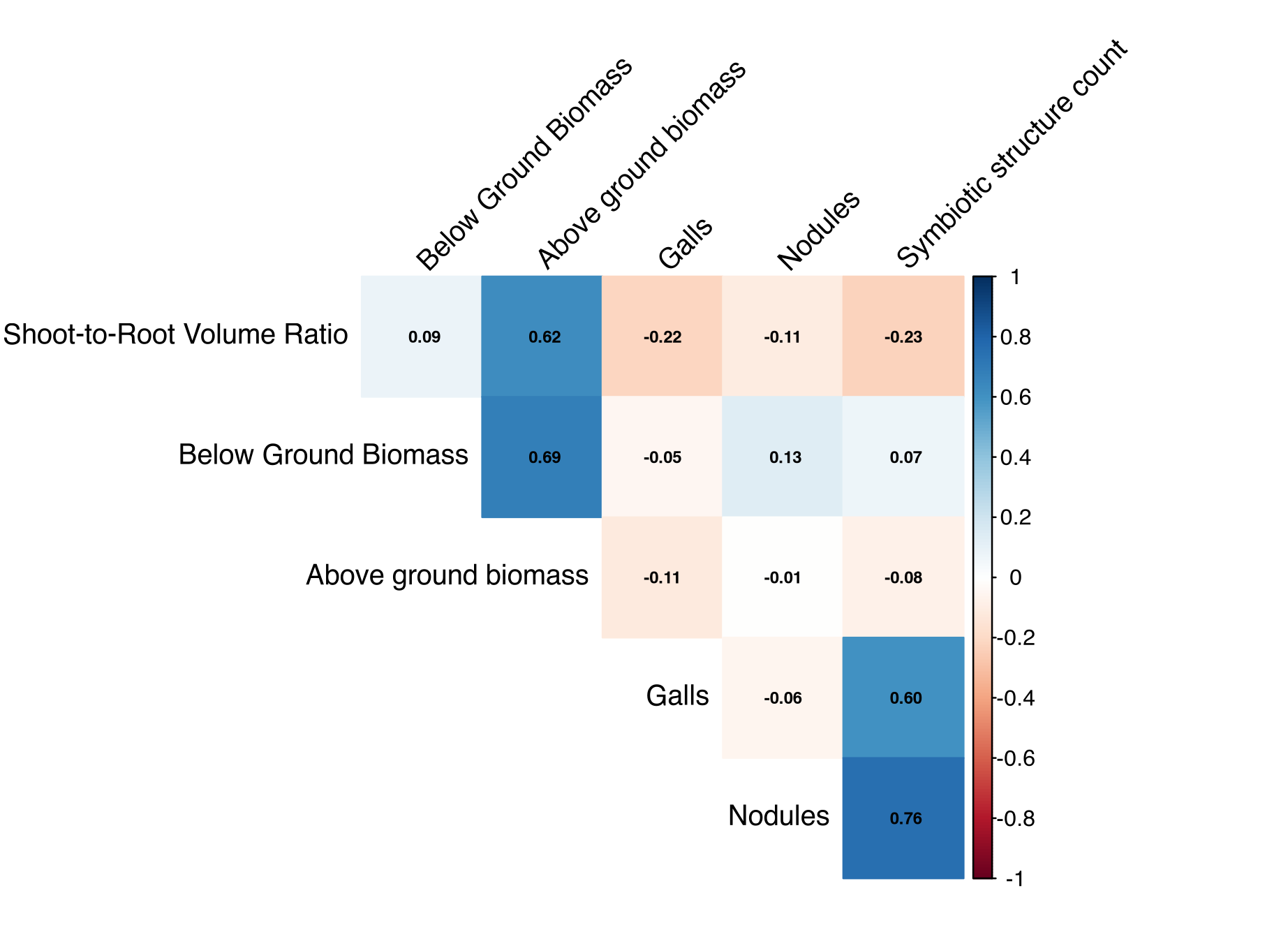

**Figure S5:** Host resistance variance component analysis that does not correct for root biomass.

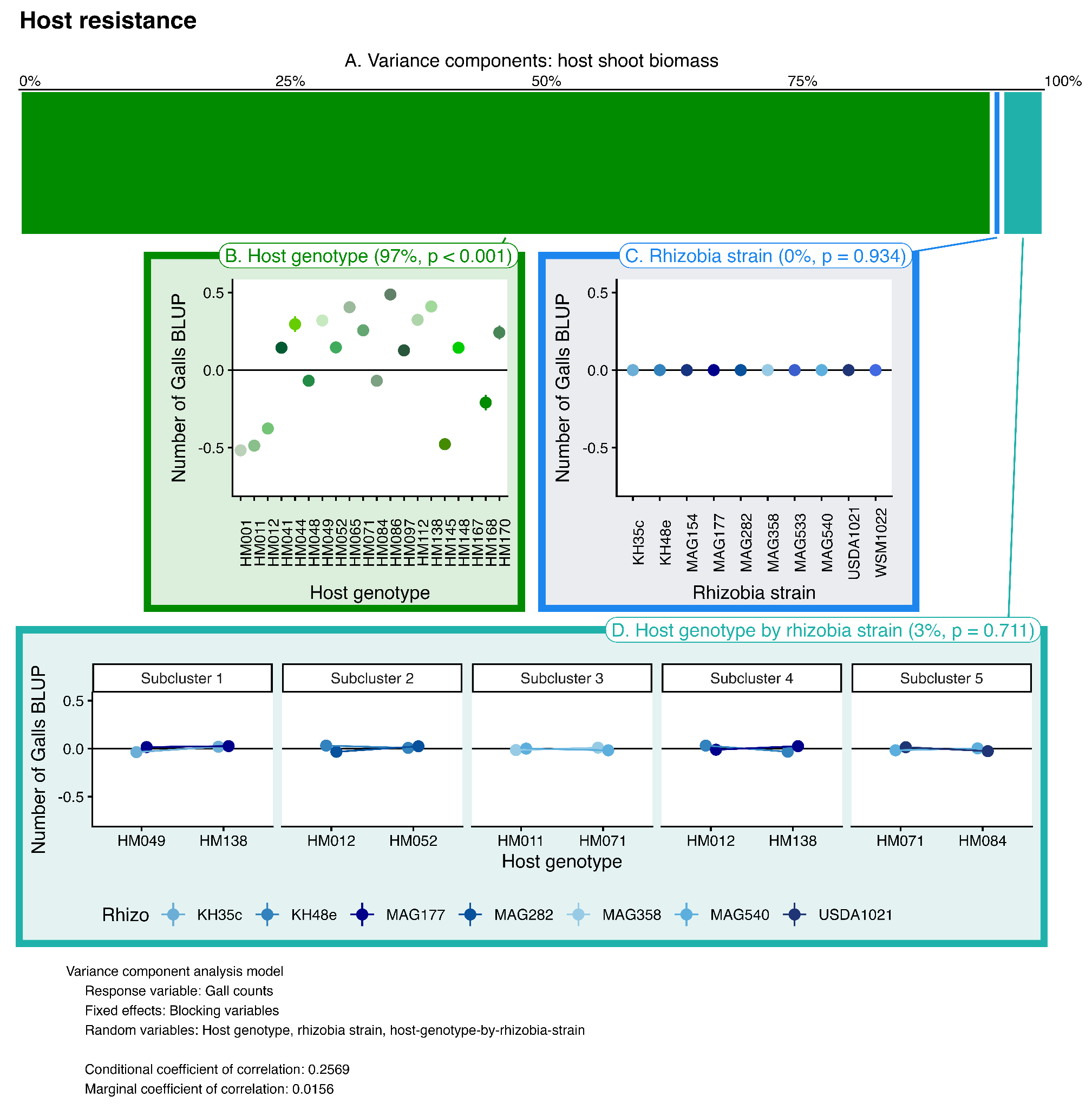

**Figure S6:** Host tolerance variance component analysis that does not correct for root biomass.

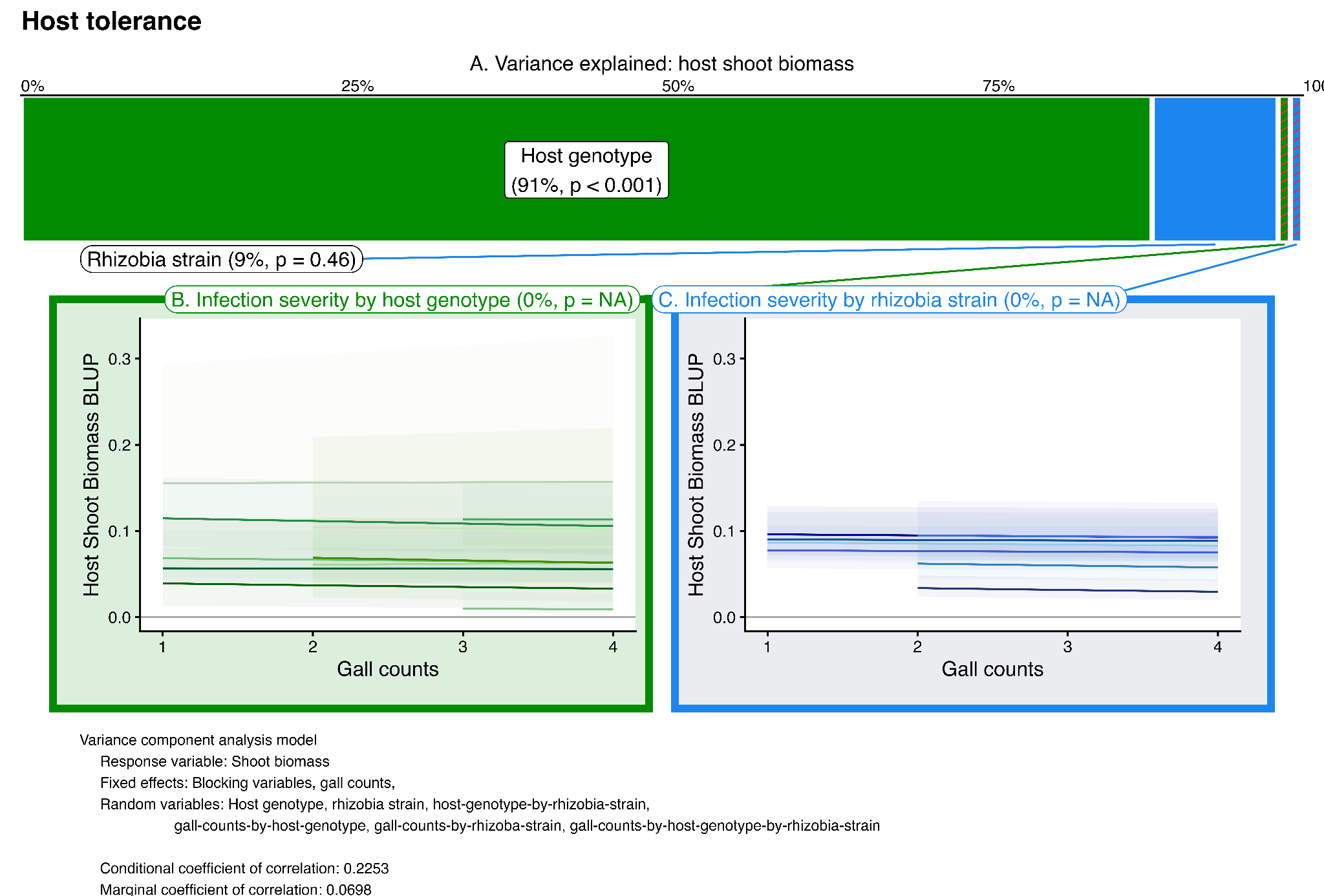

**Figure S7:** Mutualism robustness variance component analysis that does not correct for root biomass.

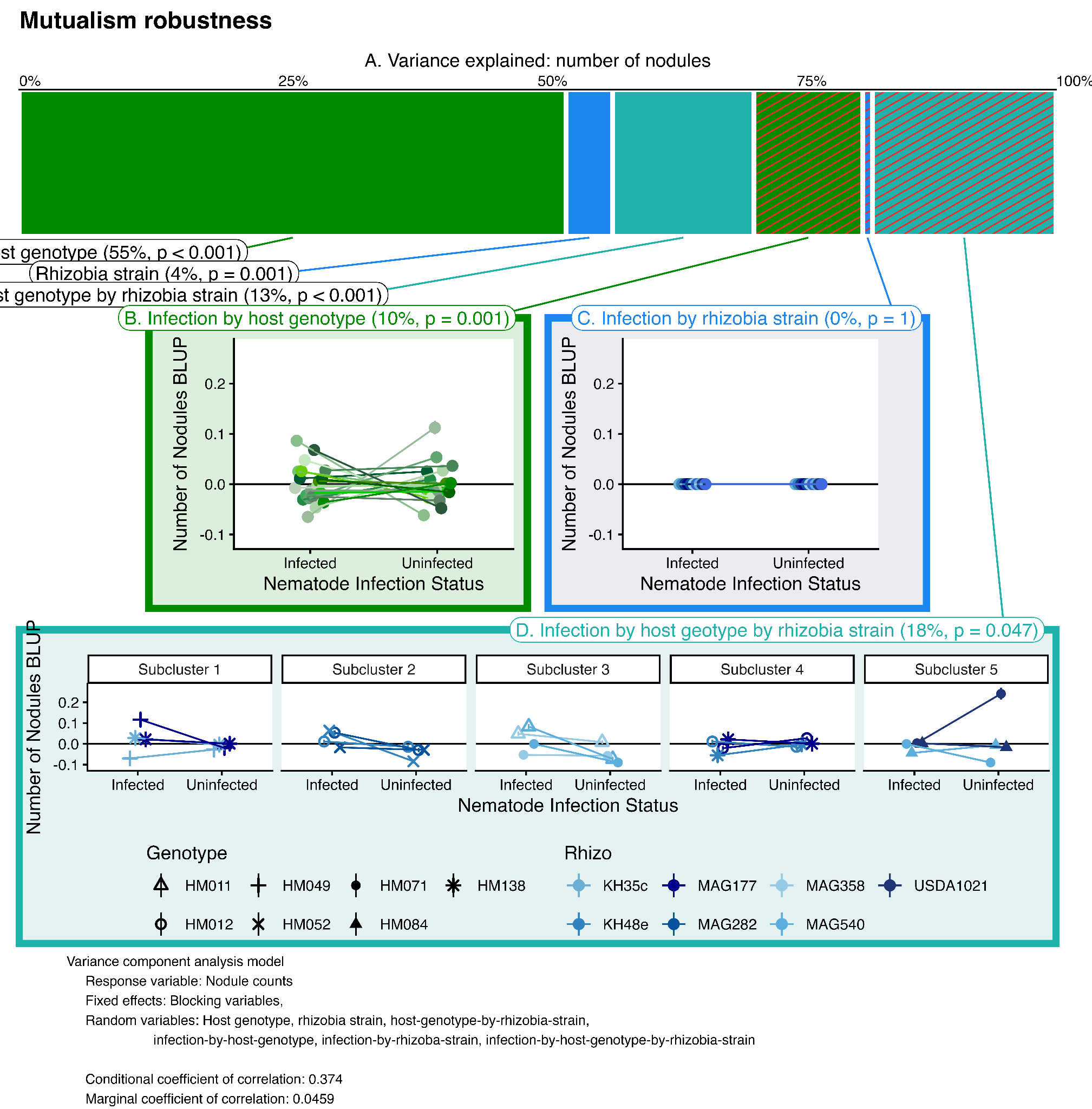
